## Supplementary Information for "Altered visual cortex excitability in premenstrual dysphoric disorder: evidence from magnetoencephalographic gamma oscillations and perceptual suppression"

* Corresponding author

### SUPPLEMENTARY METHODS, MATERIALS AND RESULTS

#### 1. Correlations between the levels of steroid hormones and GR parameters

**Supplementary table S1.** Partial Spearman's correlations between steroid hormones (estradiol, progesterone) and gamma response (GR) Power (log10-transformed) and Frequency adjusted for Age in the two groups of participants (control, PMDD). Correlations with p<0.05 (uncorrected for multiple comparisons) are highlighted in bold.

| ***A.*** *Estradiol, follicular phase* | | |
|---|---|---|
| Grating's motion velocity | Control group (N=27) | PMDD group (N=20) |
| | *GR power* | |
| Static, 0 °/s | r=-0.15, p=0.47 | r=-0.10, p=0.70 |
| Slow, 1.2 °/s | r=-0.17, p=0.40 | r=0.00, p=0.99 |
| Medium, 3.6 °/s | r=-0.17, p=0.41 | r=-0.12, p=0.62 |
| Fast, 6.0 °/s | r=-0.13, p=0.53 | r=-0.01, p=0.97 |
| | *GR frequency* | |
| Static, 0 °/s | r=0.29, p=0.16 | r=0.26, p=0.29 |
| Slow, 1.2 °/s | r=0.33, p=0.097 | r=0.20, p=0.41 |
| Medium, 3.6 °/s | **r=0.40, p=0.045** | r=-0.05, p=0.84 |
| Fast, 6.0 °/s | **r=0.41, p=0.037** | r=-0.05, p=0.85 |

| ***B.*** *Estradiol, luteal phase* | | |
|---|---|---|
| Grating's motion velocity | Control group (N=27) | PMDD group (N=20) |
| | *GR power* | |
| Static, 0 °/s | r=0.20, p=0.33 | r=-0.16, p=0.52 |
| Slow, 1.2 °/s | r=-0.03, p=0.88 | r=-0.09, p=0.71 |
| Medium, 3.6 °/s | r=-0.01, p=0.96 | r=-0.07, p=0.77 |
| Fast, 6.0 °/s | r=0.18, p=0.37 | r=-0.02, p=0.93 |
| | *GR frequency* | |
| Static, 0 °/s | r=0.05, p=0.82 | r=0.35, p=0.14 |
| Slow, 1.2 °/s | r=0.12, p=0.56 | r=0.32, p=0.18 |
| Medium, 3.6 °/s | r=0.05, p=0.81 | r=0.42, p=0.07 |
| Fast, 6.0 °/s | r=0.04, p=0.85 | r=0.35, p=0.15 |

| ***C.*** *Progesterone, luteal phase* | | |
|---|---|---|
| Grating’s motion velocity | Control group (N=27) | PMDD group (N=20) |
| | *GR power* | |
| Static, 0 °/s | r=-0.04, p=0.84 | r=-0.13, p=0.59 |
| Slow, 1.2 °/s | r=-0.25, p=0.21 | r=-0.05, p=0.84 |
| Medium, 3.6 °/s | r=-0.27, p=0.19 | r=-0.05, p=0.83 |
| Fast, 6.0 °/s | r=0.07, p=0.78 | r=-0.01, p=0.96 |
| | *GR frequency* | |
| Static, 0 °/s | r=-0.12, p=0.57 | r=0.19, p=0.43 |
| Slow, 1.2 °/s | r=0.00, p=0.97 | r=0.13, p=0.60 |
| Medium, 3.6 °/s | r=-0.05, p=0.82 | r=-0.03, p=0.91 |
| Fast, 6.0 °/s | r=-0.05, p=0.80 | r=0.05, p=0.84 |

#### 2. Gamma Suppression Slope (GSS)

##### 2.1. GSS: method of calculation

In the previous studies (Manyukhina et al., 2021; Orekhova et al., 2019, 2020), we estimated magnitude of gamma response (GR) suppression as a function of a change in the drift rate of high-contrast visual gratings (1.2°/s, 3.6°/s, 6.0°/s). The *Gamma Suppression Slope* (GSS) - the coefficient of regression of the weighted GR power to velocity - was calculated using the ‘fitlm’ Matlab function: fitlm (x, y, ‘y~x1–1’), where x = [1.2, 3.6, 6.0] corresponds to velocity of motion, y = [0, $Power_{Medium}/Power_{Slow}–1$, $Power_{Fast}/Power_{Slow}–1$] corresponds to GR power, and ‘y~x1–1’ sets the intercept of the regression line to zero. The resulting regression coefficient b is equal to zero in the case of a constant response power in the three experimental velocity conditions (i.e., ‘no suppression’) and is proportionally more negative in case of stronger velocity-related suppression of the GR.

##### 2.2. High correlation between GSS and GR suppression

There was a strong correlation between *GSS* and the *GR suppression* estimated in the present study as a normalized difference between the ‘slow’ and the ‘medium’ velocity condition (N=47; luteal Pearson’s r=-0.90, follicular Pearson’s r=-0.86, p’s <1e-13).

##### 2.3. GSS is not significantly different between the groups but does predict symptom severity in PMDD

Unlike GR suppression index, the GSS did not differentiate between PMDD and control participants (Student's t-test; follicular: t(45)=0.79, p=0.44; luteal: t(45)=1.58, p=0.12). However, similarly to GR suppression index, GSS correlated with the same-day PMS score in PMDD subjects during the luteal phase ($N_{PMDD}$=18, Pearson's r=0.51, p=0.03). Again, lower GR suppression, reflected by a less negative GSS, characterized PMDD women with more severe luteal PMS on the day of the investigation.

##### 2.4. Correlation between GSS and perceptual spatial suppression (SSI)

**Supplementary table S2.** Spearman's correlations between GSS and perceptual spatial suppression (SSI).

| MC phase | Control group (N=26*) | PMDD group (N=19*) | Difference between correlation coefficients** |
|---|---|---|---|
| Follicular | **r=-0.45, p=0.02** | r=-0.01, n.s. | p=0.14 |
| Luteal | r=-0.32, p=0.1 | r=0.20, n.s. | p=0.11 |

N – number of subjects; MC – menstrual cycle. Significant correlations and differences are highlighted in bold.

* One PMDD and one control participant were excluded because of the persistent illusion of reversed motion during the presentation of the large grating, which did not allow SSI to be estimated.

** Two-tailed.
